# An axis-referenced morphometric model reshaping the hippocampus following its longitudinal organization

**DOI:** 10.1101/2023.03.07.531443

**Authors:** Na Gao, Chenfei Ye, Hantao Chen, Xingyu Hao, Ting Ma

**Affiliations:** The Department of Electronic & Information Engineering, Harbin Institute of Technology (Shenzhen), Shenzhen, China; International Research Institute for Artificial Intelligence, Harbin Institute of Technology at Shenzhen, Shenzhen, China; Peng Cheng Laboratory, Shenzhen, China

**Author notes:** Corresponding author: Ting Ma. These authors contributed equally to this work (Na Gao, Chenfei Ye).

**Keywords:** Hippocampus, Longitudinal morphometry, laminar architecture, MRI, Computational anatomy

## Abstract

The anterior-posterior gradient of the human hippocampus on coronal slices is commonly referred to as the long axis, which exhibits morphological heterogeneity related to aging and neurodegeneration. However, the actual longitudinal organization follows a laminar architecture along a longitudinal trajectory that terminates not in the absolute anterior tip of the hippocampus, but rather traces the folding uncus. This study proposes an axis-referenced morphometric model (ARMM) to reshape the hippocampus based on its anatomical nature. An atlas of the hippocampus derived from large samples of 7T ex-vivo MRI and histology serves as the prototype of the model. We set up an orthogonal curvilinear coordinate system within the hippocampus, where the internal coordinate lines correspond naturally to longitudinal and transversal axes of the hippocampus through conformal mapping, providing reference when measuring structural features like thickness, width, length, etc. The inverse mapping allows the hippocampal grey matter to unfold onto a two-dimensional rectangle, facilitating point-wise morphological correspondence across individuals. To evaluate the ARMM, we perform a detailed shape representation of the hippocampus from 7T-MRI and compare it with three state-of-the-art shape models. The results show that the ARMM performs best in metrics measuring shape similarity between the reshaped hippocampus and the ground truth, particularly in head folds (Dice=0.9934). Moreover, our examination of ARMM on longitudinal 7T images revealed its ability to capture subtle structural variations due to neurodegeneration. The ARMM offers a unique anatomically motivated morphometric model of hippocampus, and sheds light on discovering new image markers for diseases associated with hippocampal damage.

**Graphic abstract:** 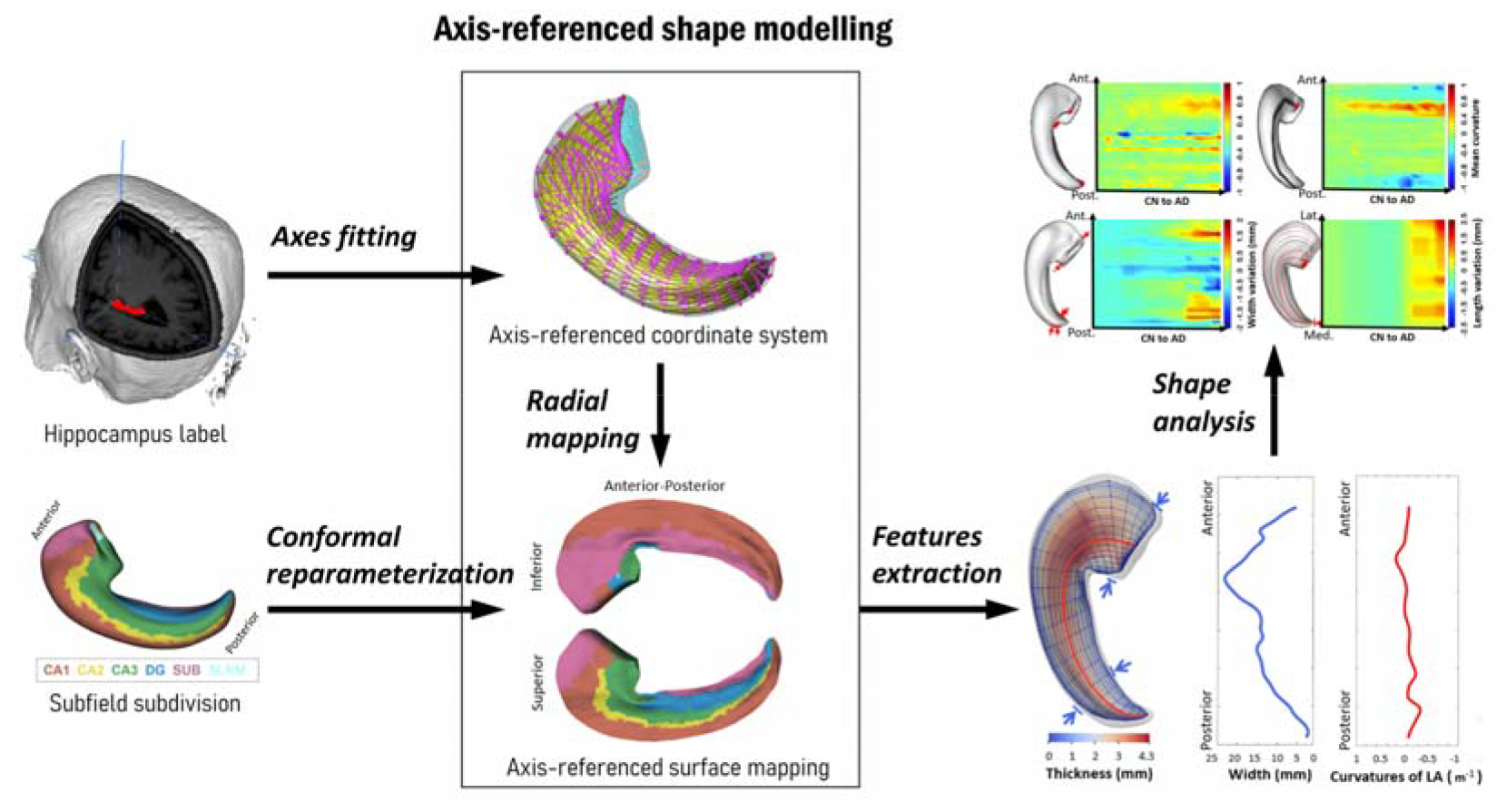

## 1. Introduction

The human hippocampus is a crucial brain structure responsible for memory processing, learning, spatial navigation and emotions (Chauhan P et al., 2021) and is vulnerable to most neurocognitive diseases (Baazaoui N et al., 2022). Recent researches have shown that the hippocampal longitudinal organization is tightly integrated into large-scale functional/structural networks (Dalton MA et al., 2022;Kharabian Masouleh S et al., 2020; Vogel JW et al., 2020). (Plachti A et al., 2020) discovers different longitudinal structural subdivisions of the hippocampus that relate to normal aging and dementias. Reconsidering the hippocampus as a heterogeneous structure along its long axis can help explain specific behavioral impairments associated with hippocampal damage. Despite the intensive interest in the longitudinal organization in MRI studies, investigating into the underpinning of this heterogeneity is difficult, as the anterior-posterior extent on coronal slices is generally considered as the longitudinal dimension of the hippocampus, ignoring its anatomical structure along the entire long axis.

The main curvature on the longitudinal dimension of the human hippocampus follows a typical curved trajectory that forms a “crescent” shape, and curves medially in the head, then superiorly, and terminates at the amygdala (Genon S et al., 2021). Although the small folds, or digitations, on the hippocampal head vary largely among individuals, this longitudinal trajectory appears to be stable. (Adler DH et al., 2018) and (Gross DW et al., 2020) demonstrate quite similar inner architecture along the entire length of the hippocampus, an “interlocking C” profile formed by the principal cell layers of CA and DG. Histological findings reveal a laminar architecture along this longitudinal curvature. Principal cell axons in the hippocampus are distributed in a parallel fashion, forming thin laminae or bands oriented nearly perpendicular to the long axis of the hippocampus. The laminae are considered to be functional units of the hippocampus. It is believed that the hippocampus processes inputs by combinations of basic laminar circuit units through interlaminar connections and that the malfunction of the laminar circuits and longitudinal connections is a root cause of aging and neurodegeneration.

Although the longitudinal organization of the hippocampus is well-described, it is challenging to represent and measure its morphology in MR images. Image-based shape analysis models, such as SPHARM-PDM (spherical harmonic description point distribution models) and s-reps (skeletal representations), generate the longitudinal axis of hippocampus but encounter difficulty following the large curvature in the head, as it is difficult to maintain continuity of internal geometric properties under large local deformation. Also, it is challenging to avoid the fan-out laminae overlapping where the cortex folds. Moreover, due to the variability of the number of digitations, traditional registration-based methods are hard to provide stable point-wise correspondence across individuals (DeKraker J et al., 2021). The unfolding method described in (DeKraker J et al., 2018) proposes a longitudinal laminar representation of hippocampal anatomy and a surface-based coordinate system for cross-individual correspondence, while not setting the laminae to be perpendicular to the long axis and requiring large manual labor for each individual hippocampus. Given a large amount of MR scans of the hippocampus in clinical practice, a computational anatomical model with a three-dimensional longitudinal organization of the hippocampus is urgently needed.

In this paper, we examine the topological structure of a hippocampal atlas derived from large samples of 7T ex-vivo MRI and histology to develop a three-dimensional computational model of the hippocampus following its longitudinal organization. The backbone of our method is to set up an orthogonal curvilinear coordinate system inside the hippocampus according to the laminar organization. Then, conformal mapping enables surface textures to transport from boundary surfaces onto medial surfaces or planar coordinates, which helps to build correspondence across individuals. Structural features that we aim to specify include the location, curvature and length of the long axis, local thickness and width along the longitudinal and transversal axes, the subfield distributions and surface curvatures. This methodological framework forms what we call the axis-referenced morphometric model (ARMM) of the hippocampus. The ARMM reshapes the uniform-distributed voxel/vertex-based hippocampal image into a laminar-distributed surface along the entire long axis. The model is verified precisely measuring the longitudinal structure of the hippocampus and able to capture subtle shape variations due to the neurodegeneration. The ARMM offers an anatomical-motivated morphological model that has the potential to generate new image markers for disease-specific hippocampal damages. All code for this project, including the ARMM template, is available at https://github.com/calliegao/ARMM.

The main contributions of this article are:

i. We characterize the laminar architecture along the entire long-axis of hippocampus using a 7T ex-vivo MRI atlas, based on which longitudinal structural features of hippocampus and the subfields can be precisely measured.
ii. We propose a uniform intrinsic coordinate system by conformally unfolding the gray matter onto two-dimensional plane, allowing for stable alignment of variable individual hippocampi.
iii. We provide an atlas template that delineates the longitudinal organization of hippocampus, which can be utilized for a wide range of clinical MRI applications, such as investigating aging or the diseases associated with hippocampal damages.

## 2. Methods

An atlas of the hippocampus, derived from large samples of 7T ex-vivo MRI and histology, serves as the prototype for our method. Initially, we generate the geometric symmetry of the hippocampus by representing it as an inscribed medial surface (IMS) inside the hippocampus. Next, we reparameterize the IMS by longitudinal and transversal orthogonal coordinate lines referencing the laminar architecture. This curvilinear orthogonal coordinate system is used to reshape the boundary surface of the hippocampus through a two-step method. Firstly, we reconstruct the implied boundary surface under medial-axis geometric constraints. Then, we refine the surfaces using conformal mapping. To validate the method’s effectiveness, we measure various geometric similarity metrics for a detailed shape representation of a 7T hippocampus. Additionally, we further validate the method on longitudinal 7T images through a strategy to automatically generate axis-referenced representations on a longitudinal series of hippocampi. Figure 1 outlines the framework of the method, and the following paragraphs provide further details.

**Figure 1.**
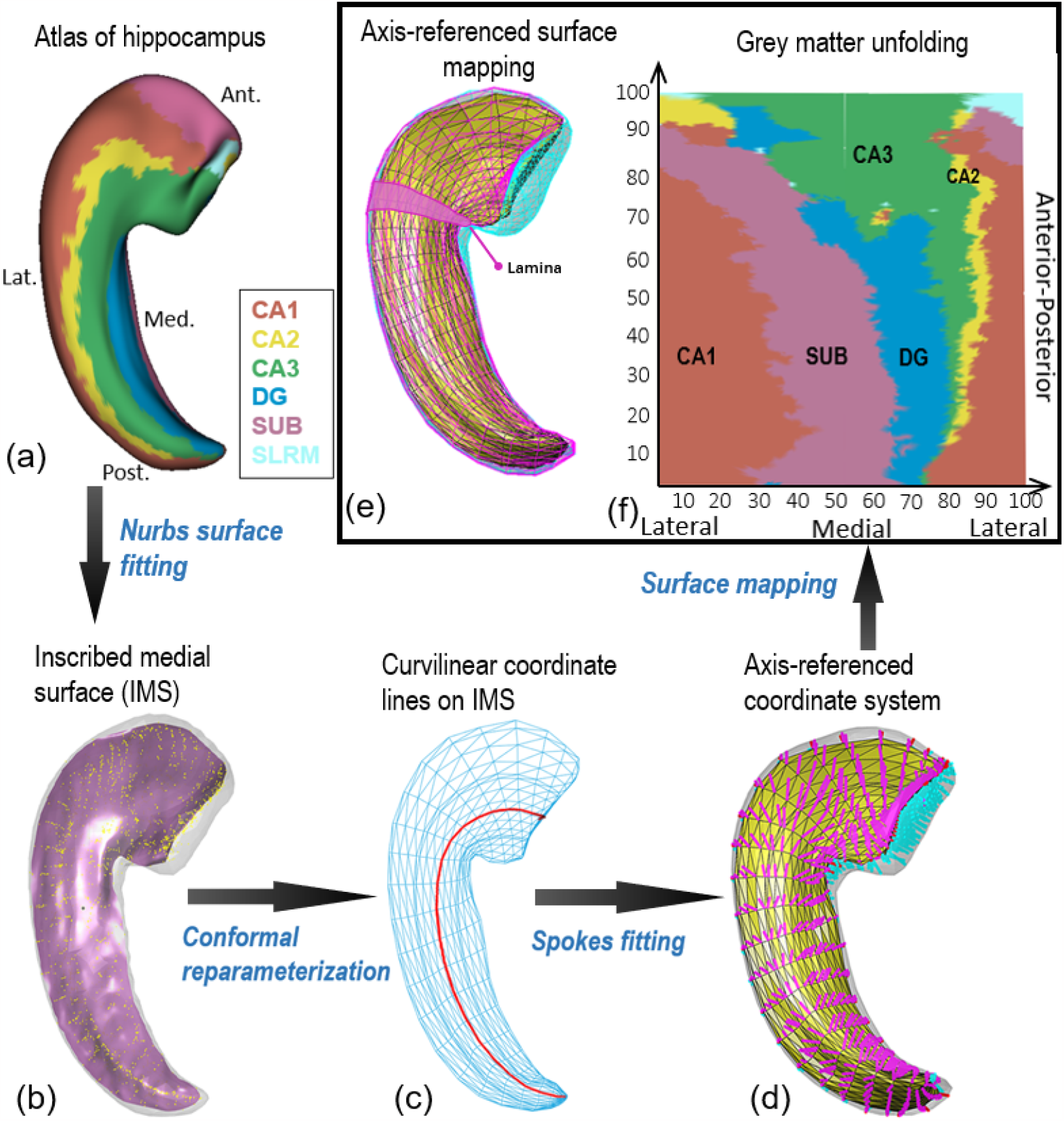
The method framework of ARMM. A hippocampal boundary surface (a) is first extracted from MR image. b) The surface is then used to calculate the IMS (pink) by Nurbs surface fitting respecting to Vironoi graph points (yellow) that generate from boundary surface mesh. Blue mesh in (c) is a reparameterized IMS by manually fitting the central symmetric axis to be agree with the CA-DG border of the hippocampus. Spokes are calculated according to geometric constrains on IMS and boundary surface. In (d), the superior and inferior spokes, colored in magenta and cyan respectively, pointing from the yellow IMS to the gray boundary surface. Finally, by a surface reconstruction method, a hippocampal surface with transversal laminar and longitudinal axes is generated from the IMS and spokes, shown in (e). Note that the boundary surface is composed of the superior and inferior parts colored in magenta and cyan respectively. Abbreviations: CA, cornu ammonis; DG, dentate gyrus; SUB, subiculum; SRLM, stratum radiatum lacunosum moleculare; Ant, anterior; Post, posterior; Lat, lateral; Med, medial.

### Dataset

We employ a probabilistic atlas of left hippocampus derived from a large cohort of ex vivo 7T MRI dataset (Adler DH et al., 2018). The data depository locates at https://www.nitrc.org/projects/pennhippoatlas/. The voxel size of the image is 0.2mm × 0.2mm × 0.2mm. The atlas includes 6 hippocampal subfields: cornu ammonis (CA) 1, 2 and 3, stratum radiatum lacunosum moleculare (SRLM), dentate gyrus (DG) and the subiculum (SUB). A longitudinal series of a left hippocampus related to progression from normal subjects to Alzheimer’s Disease (AD) patients are used in this article to verify the performance of the ARMM when characterizing disease-related subtle structural variations. The image series are also from (Adler DH et al., 2018), including 21 images (voxel size: 0.2mm × 0.2mm × 0.2mm). We extract raw surfaces of the hippocampi by ITK-SNAP (Yushkevich PA et al., 2006).

### Symmetry of hippocampal shape

The skeleton is kind of geometric symmetry to simplify a complex shape. Mathematically, the symmetry axis of hippocampus can be represented as an internal medial surface by the medial axis geometry. We make some modifications of the medial axis for applications in this study, where the details are described in supplementary material. A main modification is to introduce an inscribed medial surface (IMS) to represent the symmetry of the object. Here, we generate the IMS of the template hippocampus by three steps. First, calculate the Vironoi graph according to the boundary surface of the hippocampus (Figure 1(a)). Second, fit a NURBS surface by Vironoi points inside the hippocampus. This surface is an approximate medial surface of hippocampal shape. Finally, smoothly connect the medial surface with the boundary surface to get an IMS (the pink surface inside the hippocampal boundary surface in Figure 1(b)).

### Longitudinal and transversal axes on IMS

We represent the trajectory of the longitudinal curvature of hippocampus by a curve restricted on the IMS. Orthogonal lines on a 2D rectangle are conformally mapped to the IMS, where a central longitudinal line is manually adjusted to be best approximate to the intersection between the IMS and the CA-DG border (the CA region includes subfield CA1,2,3 and the DG includes the SRLM). Due to the conformal mapping, the transversal (medial-lateral) axes are ensured to be vertical to longitudinal axes on IMS. These orthogonal curvilinear coordinate lines then serve as a reference to reshape the hippocampal boundary surface. The conformal mapping is performed by methods described in (Meng T W, et al., 2016). We manually define four landmark vertexes on border of IMS to correspond to four corners of a two-dimensional rectangle. A reparameterized IMS is illustrated in Figure 1(c).

### Axis-referenced coordinates of hippocampus

A stratified structure implied from medial surface via radial flow provides a feasible way to index any position in the hippocampus based on the coordinate system on IMS. When specifying the target of the radial flow to be on the boundary surface, the radial mapping can be represented as a vector field. These vectors are called “spokes” in medial axis geometry. Let *S* denotes the original surface boundary of the target hippocampus, and 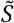 be the reconstructed surface. Spokes on IMS should satisfy the following constraints: (1) 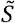 should be agree with *S*; (2) spokes are vertical to the boundary surface; (3) the lengths of superior spokes are equal to the corresponding inferior spokes. We initialize the spokes on IMS by unit normal vectors on IMS. Then, we use an iterative method to refine the initial spokes. First, to ensure the accuracy of reconstructed shape, we need the condition (1) to be strictly hold. Second, we set proper threshold measuring the non-orthogonality of spokes to boundary surface, i.e., 1 − cos (*θ*_*p*_), where *θ*_*p*_ denotes the angle between the spoke direction at vertex *p* on IMS and the normal vector of boundary surface at the nearest vertex to the spoke tips. Third, because that the spokes at bends of head often cross the boundary surface, pointing to wrong places, we solve the problem by the enforcing condition (3), then gradually adjust these spokes directions to be approximately orthogonal to the boundary surface. Finally, iterate above three steps, and take once more the implementation of condition (1) to get the final refined spokes. A Figure 1(d) shows superior and inferior spokes attached on IMS of the hippocampus.

### Reshaping the boundary surface

Theoretically, the hippocampal boundary surface can be reconstructed by smoothly connecting all spoke tips, and the laminar structure on reconstructed surface will be kept by radial map. However, in digital shape models, it is hard to satisfy all these geometric conditions. The self-overlaps of spokes will occur at large curvatures. The problem leads to self-intersecting faces on the reconstructed surface mesh. To refine the surface representations, the superior and inferior surfaces are first roughly reconstructed by superior and inferior spokes respectively. The two triangular surface meshes are subdivided automatically by the Loop algorithm. Then, each of the surface is conformally reparameterized while fixing the same landmarks with IMS reparameterization. Finally, the nearest transversal circles by conformal mapping to spokes-referenced “circles” are selected to be “borders” of “laminae”. The refined boundary surface mesh of the hippocampus is shown in Figure 1(e). A lamina is shown in the figure. The thicknesses of the laminae can be manually designed by adjusting the reparameterized mesh density. So far, we build a curvilinear orthogonal coordinate system by conformally mapping the planar coordinates onto the IMS within the hippocampus. Any vertexes on hippocampal surface can be indexed in this coordinate system.

### Unfolding the hippocampal grey matter

Since the conformal mapping builds correspondence among the IMS, the superior and inferior surfaces, and a rectangle on two-dimensional plane, subfield distributions and local curvatures on hippocampal boundary surface can be projected onto the IMS or planar rectangle through conformal correspondence. This process equals to unfold the hippocampal surface to the two-dimensional plane while remaining the geometric properties of surface textures. When inspecting topological structures of the subfields, it is known that the SRLM on slices forms a typical “C shape” inside the hippocampus, with subfields CA1, 2, 3, SUB and part of DG cutting off the external dorsal part of “C” into five parts. The internal ventral part of the “C” consists DG and part of CA3. Therefore, the unfolding of hippocampal surface keeps topological distribution of subfields that distributes around the external dorsal part of SRLM, including all the CA1, 2, SUB, most part of CA3, and part of DG. Therefore, any points on these subfields can be uniquely indexed and identified on the two-dimensional orthogonal coordinate system. An unfolded hippocampus is presented in Figure 1(f).

### Structural features extraction

Surface features, such as curvatures on each vertex, have been widely used and proved sensitive to local shape changes. Local surface features can be easily calculated on each vertex. Since the IMS and hippocampal surfaces are reparameterized follow the consistent rule that along transversal and longitudinal dimension of the hippocampus, subfield borders on hippocampal surface can be projected onto the IMS by conformal correspondence. Volumetric features, such as thickness, width and bending, have been also shown useful in disease-affected focal location. In this study, distances from mesh vertexes to the IMS are defined as local thicknesses. Since the hippocampal surface is split into superior and inferior surface along the fold curve, vertexes on each of them has respective local superior thicknesses and inferior thicknesses. This definition has several advantages: 1) Any vertexes on the hippocampal surface, including those have no corresponding spokes, have measurable thickness; 2) The thickness between the superior and inferior surfaces is straight-line/Euclidean distance, which is more easily interpreted in terms of a sheet-like shape in clinical settings; 3) Considering thicknesses of superior and inferior portions of the object respectively makes it convenient to study local shape change on each portion. Other structural features and the corresponding descriptions are listed in Table 1.

**Table 1.** The structural features and how to measure them on ARMM of hippocampus.

| *Features* | *Descriptions* |
| --- | --- |
| Surface curvatures | Mean curvatures measured on each vertex on the reshaped hippocampal boundary surface. |
| Superior thicknesses | Euclidean distances from superior boundary surface vertexes to the IMS. |
| Inferior thicknesses | Euclidean distances from inferior boundary surface vertexes to the IMS. |
| Medial-lateral widths | Lengths of transversal axes on IMS. |
| Laminar widths | Curvilinear distance between two adjacent transversal axes on IMS. |
| Lengths of the entire hippocampus | Length of the longitudinal axes on IMS. |
| Length of the long axis | Length of the long axis on IMS. |
| Curvatures of the long axis | The discrete mean curvatures measured on the long axis. |

### Model evaluation

The above methods implemented on the hippocampal atlas forms a what we called the axis-referenced morphometric model (ARRM) of hippocampus. The raw hippocampal surface extracted from the label image is used as the ground truth to test geometric accuracy of ARMM. The hippocampal surface is reconstructed by four shape models (ARMM, SPHARM-PDM (Styner M et al., 2006), ds-rep (Liu Z et al., 2021) and cm-rep (Yushkevich PA, 2009) respectively. We test four metrics that measures accuracy of the reconstructed hippocampi by the shape models, including the surface distance, areal difference, curvedness error and dice index. The surface distance Q is defined as 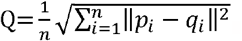, where *n* is the number of vertexes on surfaces, and *p*_*i*_, *q*_*i*_ represent the corresponding points on two surfaces. The Euclidean distance *d*_*i*_ =∥*p*_*i*_ − *q*_*i*_∥ (*i*= 1, … *n*) denotes each point-wise distance. The correspondence related to point-wise distance is constructed by searching for the nearest point between surfaces via the Iterative Closest Point (ICP) method. The curvedness at each vertex is defined by 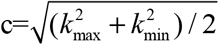, where *k*_max_ and *k*_min_ represent the main curvature. Further, we convert the reconstructed surfaces into binary image labels and rigidly register the labels into the original image label of the template hippocampus, then calculate dice index between the reconstructed shape and the ground truth. The dice index is defined as 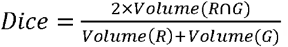, where R denotes the image label of the reconstructed shape, and G denotes the ground truth label. To evaluate the accuracy of ARMM in shape representation on different portions of hippocampus, we subdivide the hippocampus into head, body and tail by 1/3, 2/3 of the anterior-posterior extent. We cut 1/2 of hippocampal head tip along medial-lateral orientation, where consists a fold that curves medially and superiorly. We further cut 1/2 of the hippocampal tail tip along the anterior-posterior orientation.

To automatically generate ARMMs of individual hippocampi, we transform the ARMM of the template model to the target hippocampus by a diffeomorphic transformation method provided by the deformetrica toolkit (Fishbaugh J et al., 2017). We employ a longitudinal sequence of 7T MR hippocampal images, which describes the disease progression from cognitive normal (CN) persons to Alzheimer’s disease (AD) patients. The image series include 21 images. For simplicity, we let each of the images denote a single time point. To generate ARMMs for a time series hippocampi, first, we rigidly align all the hippocampi to the baseline hippocampus. Second, we fit the baseline hippocampus with a ARMM by the deformation strategy. Third, hippocampi on each other time points share the same IMS of the baseline hippocampus, while optimize the spokes and fold curves to fit each hippocampus boundary surfaces. Finally, the boundary surface on each time point is reconstructed by using the surface reshaping method described in above paragraphs. Local surface variations are quantified by calculating point-wise surface distance between the baseline and the *i*^*th*^ observed surface. The point-wise surfaces correspondence is established by coordinate system of ARMMs. Thickness variation is evaluated by local superior/inferior thickness measured on each vertex of the reshaped hippocampal surface. Since the AD-related shape variations are dominated by atrophy, all variation measurements are calculated by subtracting features of the observed hippocampi from the features of the baseline hippocampus.

## 3. Results

### Model accuracy on the template hippocampus

We applied the ARMM to an atlas of hippocampus obtained from 7T MRI and compared its accuracy to that of three other shape models using surface distance, areal difference, curvedness error and dice index. The results are presented in Table 2, which shows that the ARMM outperformed the other models in all portions of the hippocampus, particularly in the head fold and tail tip, where it shows high accuracy. The areal difference of ARMM is found to be higher than that of SPHARM-PDM, which is expected, given the relatively sparse point distribution on the lateral surface of the hippocampus due to the fan-out laminae from the lateral to medial. The ARMM also performed well on portions of the hippocampus with large curvature, as indicated by its good curvedness and surface area.

**Table 2.** Shape errors between each reconstructed shape and the ground truth.

| *Portions of hippocampus* | *Shape similarity metrics* | **ARMM** | Cm-rep | SPAHRM | Ds-rep |
| --- | --- | --- | --- | --- | --- |
| *Whole hippocampus* | *Surface distance (mm)* | **0.0003** | 0.0071 | 0.0012 | 0.026 |
|  | *Areal difference (mm<sup>2</sup>)* | 1.2404 | **0.4222** | 1.7255 | 9.9669 |
|  | *Curvedness error (1/mm)* | **0.0133** | 5.0635 | 0.0326 | 0.0592 |
|  | *Dice* | **0.9961** | 0.8313 | 0.9702 | 0.9074 |
| *Head* | *Surface distance (mm)* | **0.0002** | 0.0054 | 0.0009 | 0.0028 |
|  | *Areal difference (mm<sup>2</sup>)* | **0.5217** | 7.4814 | 1.5972 | 11.3964 |
|  | *Curvedness error (1/mm)* | **0.002** | 6.9358 | 0.0293 | 0.0643 |
|  | *Dice* | **0.9959** | 0.8277 | 0.9732 | 0.9045 |
| *Head fold* | *Surface distance (mm)* | **0.0003** | 0.0069 | 0.001 | 0.0044 |
|  | *Areal difference (mm<sup>2</sup>)* | **0.6414** | 1.2509 | 1.3612 | 17.8808 |
|  | *Curvedness error (1/mm)* | **0.0024** | 14.136 | 0.0224 | 0.0573 |
|  | *Dice* | **0.9934** | 0.6815 | 0.9706 | 0.8459 |
| *Body* | *Surface distance (mm)* | **0.0004** | 0.0106 | 0.0012 | 0.1137 |
|  | *Areal difference (mm<sup>2</sup>)* | **0.1076** | 8.1713 | 0.127 | 5.4684 |
|  | *Curvedness error (1/mm)* | 0.0679 | 0.0281 | **0.0134** | 0.0146 |
|  | *Dice* | **0.998** | 0.8403 | 0.9722 | 0.9284 |
| *Tail* | *Surface distance (mm)* | **0.0002** | 0.0046 | 0.0012 | 0.0031 |
|  | *Areal difference (mm<sup>2</sup>)* | **0.2534** | 17.5162 | 2.0229 | 10.8307 |
|  | *Curvedness error (1/mm)* | **0.0029** | 0.0095 | 0.0402 | 0.0501 |
|  | *Dice* | **0.997** | 0.8313 | 0.9611 | 0.8934 |
| *Tail tip* | *Surface distance (mm)* | **0.0004** | 0.008 | 0.0017 | 0.0061 |
|  | *Areal difference (mm<sup>2</sup>)* | **0.3948** | 28.8729 | 4.0824 | 25.5454 |
|  | *Curvedness error (1/mm)* | **0.0036** | 0.0445 | 0.0369 | 0.0037 |
|  | *Dice* | **0.9939** | 0.7272 | 0.9517 | 0.7747 |

Figure 2A visualizes the local surface distance between the reconstructed surfaces and the ground truth. The SPHARM-PDM and ARMM performed well in characterizing the hippocampal head and tail, while the ds-rep and the cm-rep models fail to capture the bending on these portions. The ARMM shows higher accuracy than SPHARM, particularly at the bending on the medial side of the head. The laminae on the hippocampal surface are generated from ARMM and are visualized as black curves in Figure 2A, showing a fanned-out distribution from the lateral to medial of the hippocampus, without any overlap at the folds.

**Figure 2.**
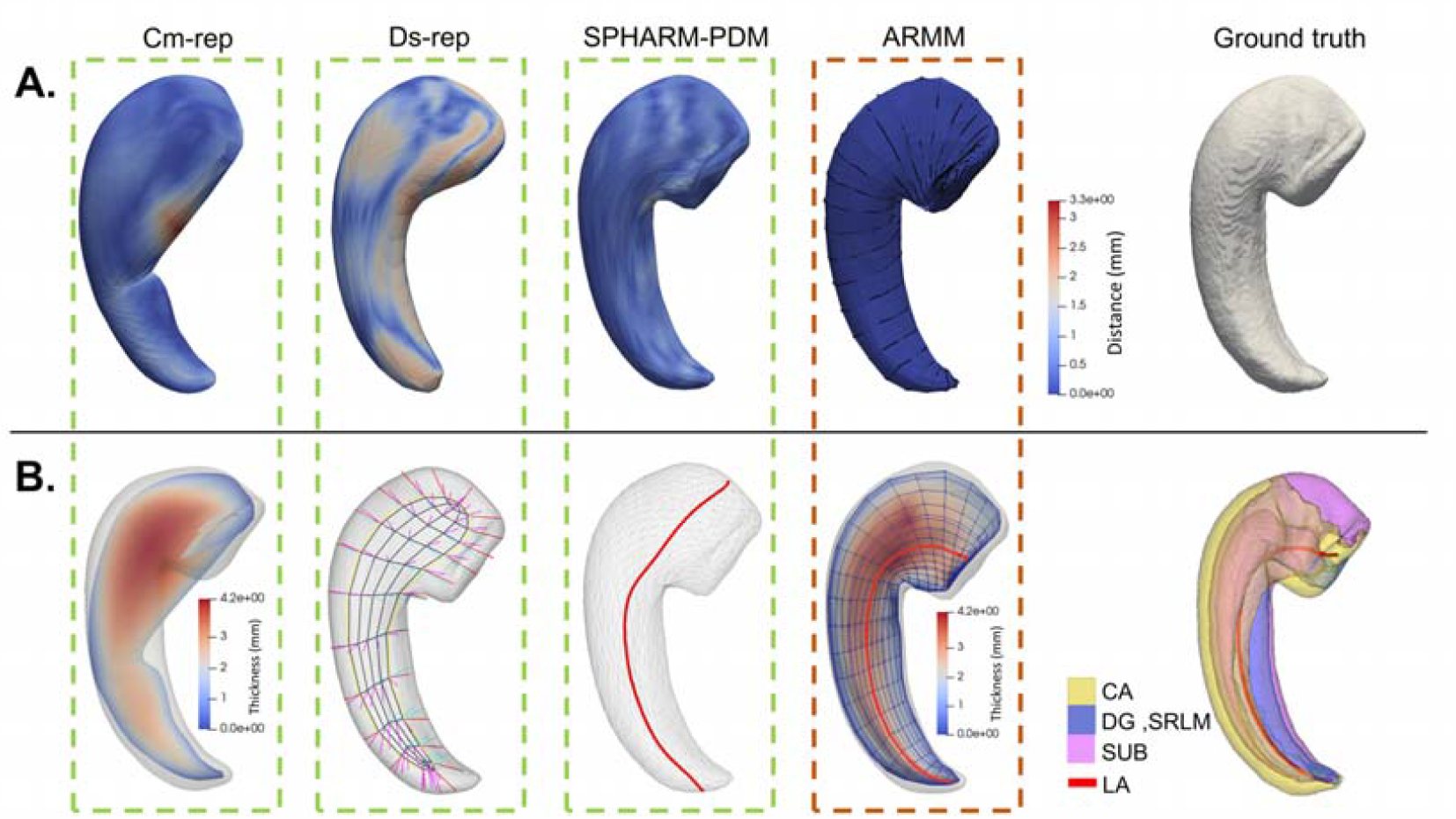
Comparisons of the four shape models that fit the same hippocampus. A) Point-wise distance maps of the models. The locations colored in red denote large errors between the ground truth surface and the reconstructed surface by the model. B) Axes that generate from these models. In the last panel, the long axis generated from ARMM is visualized inside the 3D hippocampus. Abbreviations: CA, cornu ammonis; DG, dentate gyrus; Sub, subiculum; LA, long axis.

Figure 2B shows the long axes of the hippocampus from the four models. The ds-rep and SPHARM-PDM well characterize the curved body of the hippocampus but failed to capture the bending on the hippocampal head and the end of tail. The cm-rep cannot generate a longitudinal axis directly from its medial surface. None of the three models consider interior anatomical structure of the hippocampus. The last panel of Figure 2B shows a 3D visualization of the subfields distribution and the long axis derived from ARMM, which agrees with the CA-DG border and follows the orientation of the entire longitudinal curvature.

The cm-rep and ARMM in Figure 2B show local thickness map measured on the parametrized IMS. The curved transverse and longitudinal coordinate lines are nearly orthogonal to each other on IMS. We also calculate the angular distortion of conformal reparameterizations of the superior surface, inferior surface and IMS, where the angular distortion refers to the absolute value of the difference between angles of the triangle elements on the 2D rectangle and corresponding angles on the conformal reparameterization. Almost all of the angular distortions are found to be less than one degree, ensuring that the laminae are perpendicular to the longitudinal axes.

### Ability to characterize disease-affected shape variations

The features characterizing the shape variations of the hippocampus during the progression of AD are presented in Figure 3. Local surface and thickness variations at four time points (the 15th, 17th, 19th and 21st time points) are compared to the baseline (the 1st time point) hippocampus. The results are displayed in Figure 3(a) and Figure 3(b), which show more severe atrophy on the inferior than the superior during disease progression. The primary atrophy occurs on the inferior lateral of the body, while focal atrophy is distributed on the head and near the endpoint of tail. The thickness variations (depicted in Figure 3(b)) are consistent with the surface distance variations. The thickness decreases are diffused and focally distributed along the long axis of the hippocampus, gradually spreading from the middle portion of the head to the posterior during AD progression.

**Figure 3.**
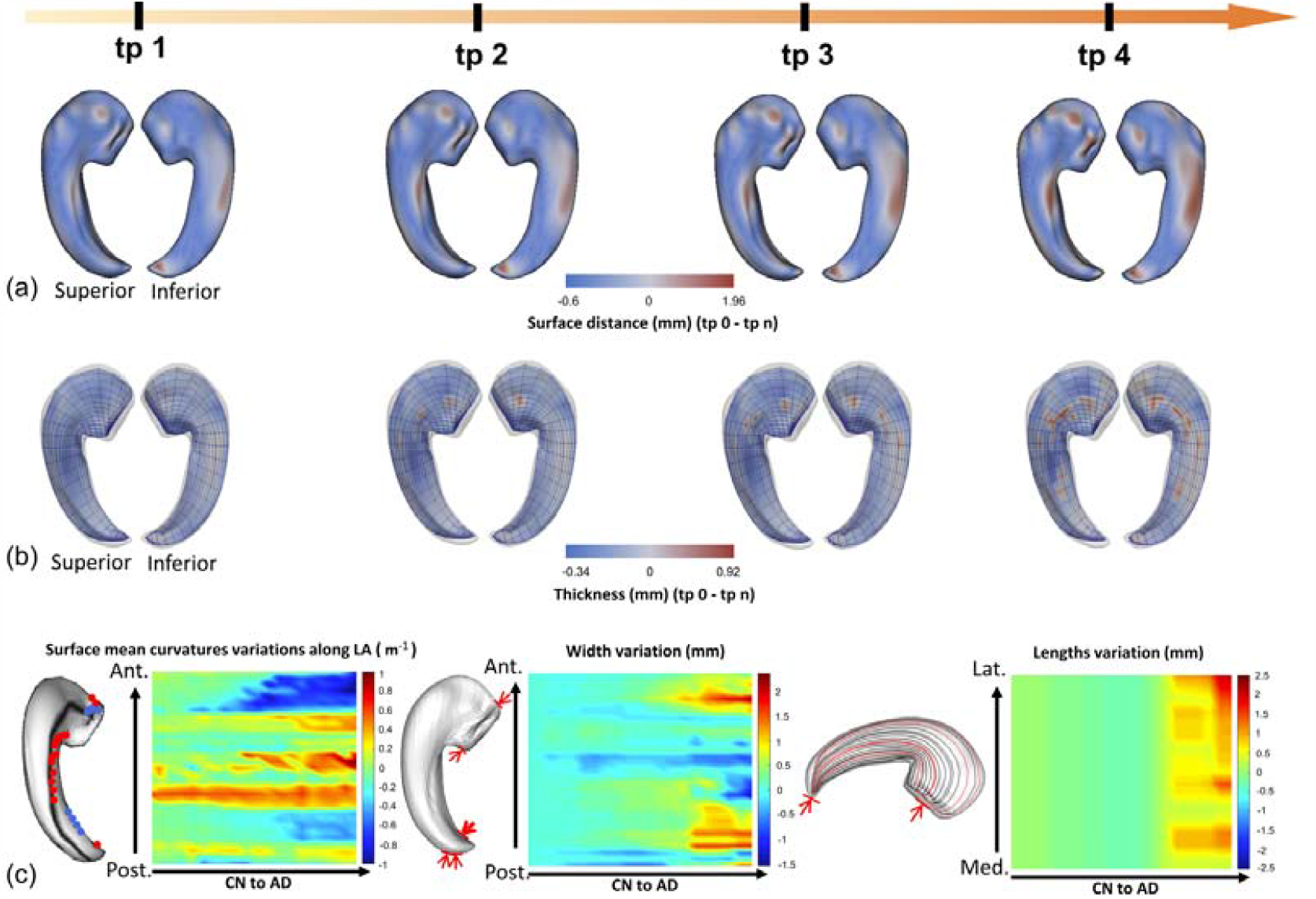
Morphological features calculated from ARMM that measure local shape variations on a time series of left hippocampi. a) Surface local variations on the superior and inferior of hippocampus at four selected time points. The focal atrophy is denoted by surface contraction (red). b) Superior and Inferior thickness local variations at four selected time points, where the thickness decrease is colored in red. c) Typical local surface curvature variations along a longitudinal axis. Curvature decreases/straightening (red); curvature increases/bending (blue). d) Hippocampal lengths variation along transverse axes during AD progression. Note that the color red in all colorbar denotes decreases of the measurements compared with baseline, while blue denotes the increase of the measurements. Abbreviations: tp, timepoint; LA, long axis; CN, cognitive normal; AD, Alzheimer’s disease.

In the left panel of Figure 3(c), the typical locations on the hippocampus that suffer relatively large curvature variations during disease progression are shown. The red points denote where the mean curvature decreases, and blue points denote where the mean curvature increases. These points are distributed near a specific long axis, the black curve shown on the hippocampal surface. The right panel of Figure 3(c) shows curvature variations of these points along the longitudinal axis on the 20 observed hippocampi from CN to AD. At the locations where the surface curvature is positive (convex), such as the middle superior of the body and the top of head, curvature tends to decrease over time. Conversely, at locations where the surface curvature of the surface is negative (concave), such as the superior of the bending of head and the lateral tail, curvature tends to increase. This result reflects of local atrophy on the hippocampus.

Figure 3(d) shows the width variations respecting the entire length of the hippocampus during disease progression. The left panel of Figure 3(d) explains how we measure the widths of the hippocampus along the longitudinal curvature of hippocampus, and the red arrows point to locations where large width variations occur. The right panel of Figure 3(d) shows width variations along the long axis of entire hippocampus across the CN to AD progression. Width decreases occur mainly at the portion near the top of the head and end of tail, while the width near the first bending of the head increases. Figure 3(d) also illustrates the entire hippocampal length variations. The left panel explains how we measure these lengths, and the red curves are parametric lines that suffer relatively large variations over disease progression. The diagram on the right shows the lengths variations during CN to AD progression. We find that the lateral of the hippocampus has the greatest length decreases.

## 4. Discussion

We examine the topological structure of a hippocampal anatomical atlas and use an axis-referenced morphometric model (ARMM) to capture the longitudinal organization of the hippocampus. The ARMM provides an intrinsic coordinate system that reshapes the hippocampus and allows for the measurement of structural features along the axes and laminae of the hippocampus. We verify that the ARMM accurately captures the longitudinal hippocampal formation and shape variations caused by neurodegeneration. When using this model, there are three aspects of the ARMM that need to be considered: (1) the potential to represent hippocampi using this anatomically motivated model on clinical data, (2) the accuracy of the laminar structure along the explicit long axis by the ARMM for the hippocampus, and (3) the stability of the point-wise correspondence provided by the ARMM coordinate system. We will discuss these three aspects as well as the limitations of the method.

### Clinical application of ARMM: from anatomical atlas to 3T in-vivo MRI measures

Increasing studies suggest that specific regional vulnerability within the hippocampus differentiates AD, vascular disease, aging, depression, and post-traumatic stress disorder (Small SA et al., 2011). Some studies highlight the role of variabilities on the longitudinal organization of the hippocampus (Bernhardt BC et al., 2016;Dautricourt S et al., 2021;Decker AL et al., 2020;Kalmady SV et al., 2017;Langnes E et al., 2020;Lee JK et al., 2020;Szeszko PR et al., 2003). However, few of them consider the anatomic nature of hippocampus. The main contribution of this study is that the ARMM can represent the hippocampus in a delicate way and provide anatomical-motivated structural features. It is promising for the model to be applicated on clinical data. To preliminarily verify this potential, we propose a strategy to automatically fit ARMMs for a group of hippocampi. The key point is to use a large deformation-based method to spatially transform the template hippocampus and its ARMM coordinate system to the target hippocampus. Details of the method are presented in the Model Evaluation section. We conduct an auxiliary experiment to verify this method using 5 7T ex-vivo MRI of hippocampi, which is detailed in supplementary materials. The result shows that the ARMM has higher accuracy than ds-rep in terms of point-wise distance errors when representing the same hippocampi. The higher accuracy may result from the advantages of the deformation algorithm in dealing with large local differences, especially in the head folds. The means of surface point-wise distance error of ARMM are 0.08mm. The maximum surface point-wise distance between the ARMM and the ground truth is 0.214mm, comparable to the minimum resolution of the images, which is fairly satisfactory. Moreover, the transformation keeps the topology of the long axis that follows the main longitudinal curvature and folds of the hippocampus, shown in Figure S1 of the supplementary documentation. The results indicate that the deformation strategy allows all the elements of ARMM to be transformed into the target hippocampus from MR images while achieving a resolution-level accuracy in generation of new ARMMs. This shows great potential for applying the ARMM on extensive clinical 3T in-vivo MRI applications.

### The long axis of the hippocampus by ARMM

In this study, laminar structures on surface are determinate by transverse axes that are orthogonal to longitudinal axes, where the central longitudinal axis follows the intersection between the CA-DG border and IMS. This longitudinal axis is what we defined as the long axis of hippocampus. While currently, there reaches no consistency for the location and morphology of the long axis of hippocampus. The current large number of studies on the longitudinal organization adopt the tripartite model that subdivides the hippocampus into head, body and tail, as is indicated by the red dotted lines shown in Figure 4. Many recent studies suggest the entire length of hippocampus to be considered in morphological analysis. (Adler DH et al., 2018) take the medial curvature as the long axis, while (Gross DW et al., 2020) take the lateral curvature as the long axis. (DeKraker J et al., 2018) proposes a method to trace the entire hippocampal length through longitudinal potential field on SRLM by Laplace equation. They verify the SRLM shape can well capture the general hippocampal shape including the complex folding patterns on the head. This finding is close to the ground truth of the “long axis”. While, the paper also finds that the SRLM are often not visible in the most medial, vertical component of the uncus, where must be manually defined. Despite the many uncertainties, we can confirm that the long axis of hippocampus has its natural anterior termination in the more medial and posterior vertical component of the uncus, and all subfields of the hippocampus contiguously follow this curvature through the hippocampal head (Ding SL and Van Hoesen GW, 2015). These evidences reveal that the long axis at head is not in the absolute anterior tip of the hippocampus that defined on coronal slices.

**Figure 4.**
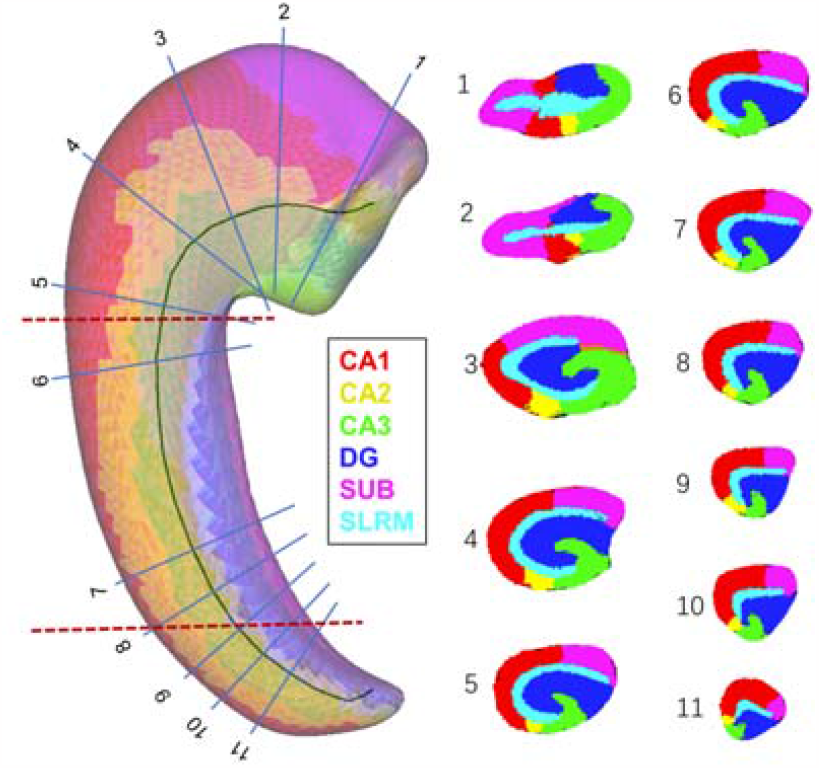
Representative slices that cut perpendicularly to the ARMM-derived long axis. Left: traditional subdivision method (red dotted lines) and slicing along directions perpendicular to the long axis (blue lines). Right: Slices corresponding to the cutting orientations shown in the left panel.

Large cross-individual differences on hippocampus should also be considered in the shape model. The different of number and scale of digitations on head and tail cause the lateral/ventral curvature fluctuated and deviated from the natural curvature of the entire hippocampus. This is also the cases for the SRLM. The variable folding patterns will lead to diverse long axes among individuals, which is disadvantage for population analysis of hippocampal morphology. Moreover, most protocols for clinical image segmentation do not segment the digitals, or do not honor its complexity and variability (Yushkevich PA et al., 2015). We should also not expect to trace the long axis by SRLM subdivision on clinical images. However, actually, the general curvature of hippocampi on the longitudinal dimension shares a consistent trajectory, for example, the left hippocampus curves like a crescent from posterior to anterior, and curves medially, posteriorly, then superiorly. This trajectory moderates the folding patterns caused by variable digitals, which is more feasible for statistical analysis.

We find that the ARMM generated long axis often locates on border between CA3 and DG, around the CA3 pyramidal cells that near the subgranular zone (SGZ) of DG (shown on slices in Figure 4), except for the portions near the head and tail endpoints. Some evidences to support reasonability of this long axis include that, first, the “CA3-centric” view of hippocampal information processing (Scharfman HE, 2007), which believes that CA3 pyramidal cells projecting diversely to all hippocampal principal cells. Second, many studies demonstrate the existence of excitatory projecting associational pathways of dentate mossy cells and CA3 pyramidal cells (Gloveli T et al., 2005;Pak S,Jang D,Lee J,Choi G,Shin H,Yang S and Yang S, 2022;Ropireddy D et al., 2011). Third, (Gross DW et al., 2020) demonstrates the internal subfield consistency inside hippocampal head. We compare the subfields on slices along the long axis of our template hippocampus with the subfield distribution on similar located slices of a typical hippocampus in the literature. In Figure 4, all slices from head to tail show the “interlocking C” or the “body-like” profile discovered in (Adler DH et al., 2018;Gross DW et al., 2020). This result shows consensus inner structures along the ARMM long axis and the manual tracing method, and further indicated the feasibility to proximate the “real” long axis by the ARMM long axis.

### Morphological correspondence by ARMM

The determination of point-wise correspondence across objects is a key subject in morphological analysis. Ambiguous correspondences at locations with large individual variabilities may lead to completely different results in statistical analysis. According to (Ding SL and Van Hoesen GW, 2015), the hippocampal head has different numbers of digitations across subjects, posing a major challenge in establishing correspondences. The variability in hippocampal digitations may not be well suited for registration-based approaches, as it is inconclusive how to align the anterior portions among populations with different numbers of digitations (DeKraker J et al., 2021). Furthermore, the correspondence established by current geometric methods is not always directly related to or supported by physiopathological evidence.

Adler discovers that the hippocampal tail (the posterior 1/3 of the hippocampus), while slicing it at approximately 6/9, 7/9, 8/9 of the total hippocampal length along the curvature of the long axis, very few samples have different subfields distribution (Adler DH et al., 2018). This finding indicates a stable correspondence between individuals along the curved long axis. However, there are limitations for inter-individual morphological correspondence by proportions along the long axis. The hippocampus may not undergo isotropic atrophy under the effect of disease. Therefore, the length and curvature of the long axis may change, especially in the head and tail of the hippocampus, leading to deviations in the correspondence. As a result, the correspondence established based on proportions is no longer valid, even if the effect may not so extreme. In (Poppenk J et al., 2020), the author describes a similar mis-correspondence example that due to the contraction of the uncus, a larger portion of anterior is redefined as posterior of hippocampus, leading to incorrect conclusion that the posterior segments grow. One possible solution is to use the long axis of the healthy hippocampus, and fit the corresponded ARMM coordinates directly to the atrophied hippocampus to the same subject, followed by eroding extra elements outside the boundary surface (i.e., using the longitudinal strategy proposed in this paper).

Evidence for long-axis-based intra-individual correspondence is rare, likely due to difficulties in sample acquisition and manual annotation. At most scenarios, shape changes on the hippocampus of the same individual have little influence on its overall shape, so the internal skeleton of the baseline observation remains stable over time. In section 3.2, we measured local shape variations on time series of hippocampi and find that the focal distribution measured by surface and volumetric features are highly consistent. This finding indirectly verifies the intra-individual correspondence by the long axis. It is highly recommended to use the ARMM to build longitudinal correspondence across hippocampi.

### Limitations

There are still some limitations to the current model. First, our method needs to separately deal with the superior and inferior surfaces of the hippocampus, which can introduce errors in identifying where the surfaces connect. A more effective approach to process the overall boundary surface will improve computational efficiency and reduce errors. Second, the proposed method does not support representing the subfield morphology within the hippocampus. To provide more structural features of the hippocampus, such as subfield shape features on laminae, it would be beneficial to include subfield morphometry in our model.

## 5. Conclusions

The conventional anterior-posterior tripartite subdivision of the hippocampus has restricted the exploration of the underlying mechanisms of the longitudinal heterogeneity of hippocampus. This paper proposes a novel framework to reshape the hippocampus based on its longitudinal organization, which allows for a detailed characterization of anatomically motivated structural features on hippocampal images. The proposed framework, called the axis-referenced morphometric model (ARMM) of the hippocampus, can represent and measure the hippocampal morphology along its entire long axis and laminae. The ARMM is verified to precisely capture the longitudinal hippocampal formation and subtle shape variations on the hippocampus due to neurodegeneration. This method has three advantages:

i. ARMM characterizes laminae (considered as functional units of hippocampus), along the entire long axis of hippocampus, providing more finely detailed anatomical information than subfields.
ii. ARMM allows for the stable alignment of hippocampi across individuals with variable morphologies by a uniform planar orthogonal coordinate system.
iii. The method offers a way to automatically generate anatomically motivated structural features on in-vivo MRI that characterizes aging and diseases associated with hippocampal damage.

Future directions for this work include the integration of subfield morphologies into this model to quantitatively analyze subfield variations on laminae. This method has the potential to assist investigations of the longitudinal heterogeneity of the hippocampus, as well as discovering new structural markers for neurodegenerative diseases.

## Acknowledgement

This study is supported by grants from the National Natural Science Foundation of P.R. China (62276081, 62106113), Innovation Team and Talents Cultivation Program of National Administration of Traditional Chinese Medicine (NO:ZYYCXTD-C-202004), Basic Research Foundation of Shenzhen Science and Technology Stable Support Program (GXWD20201230155427003-20200822115709001).

## Data and Code Availability Statement

Data supporting the findings of this study is based on opensource results of (Adler DH et al., 2018). The data depository locates at https://www.nitrc.org/projects/pennhippoatlas/. All code for this project including the ARMM template is located at https://github.com/calliegao/ARMM. The code for the pipeline is publicly available after acceptance.

