## supplementary material for "An axis-referenced morphometric model reshaping the hippocampus following its longitudinal organization"

1. **Theoretic background of ARMM: The medial axis geometry and its modification for application on computational images**

It is simple and intuitive to express an object’s general shape by its skeleton, because it expresses the basic topology of the object. Objects of the same kind tend to have same bends and branches. When we aim to describe differences between similar objects or the object’s deformations, more detailed descriptions on object boundary are necessary. Blum (Damon J et al., 2003; Damon J et al., 2008) first proposed a medial representation (m-rep) method, which can completely reconstruct the boundary from its skeleton. In other words, the method is a complete representation of the shape based on skeleton. Damon makes rigorous mathematical study of m-reps. Pizer (Pizer S et al., 2006) develops skeletal representations (s-reps) generation algorithms from given objects and statistical analysis method of based on s-reps. The method has been widely applied to medical image analysis and other many fields.

In medial representations, the medial axis of an object, also called the medial locus, is formed by trajectory of maximal inscribed balls (MIBs) inside the object. The medial axis consists of two parts: the trajectory and the maximum radius of MIBs. Formally, consider an unbranching genus zero 3D object. Let its medial manifold be $\left( m,r \right)\in R^{3}\times R^{+}$, where $m$ is a continuous smooth surface in $R^{3}$, and $r$ is a scalar field on m. Since any point on the boundary has only one corresponding MIB, the boundary surface can be completely reconstructed by MIBs. Let each point on the boundary $S$ correspond to a vector $\boldsymbol{r}$. $\boldsymbol{r}$ starts from $m$ and ends at $S$, and is perpendicular to $S$. The vector field $\boldsymbol{r}$ has three properties. Consider any point $p$ on $m$, it has two vectors $r^{+},r^{-}\in\boldsymbol{r}$. Let the normal vector of $m$ at $p$ be $n_{p}$, $r^{+}$ point from p to $S_{p}^{+}\in S$, and $r^{-}$ point from p to $S_{p}^{-}\in S$. Let the normal vectors of S at $S_{p}^{+}$ and $S_{p}^{-}$ be $n_{S_{p}^{+}}$ and $n_{S_{p}^{-}}$. Then 1) $<n_{p}{,r}^{+}>=<r^{-}{,-n}_{p}>$ , 2) $\left\| r^{+} \right\|= \left\| r^{-} \right\|$, and 3) $n_{S_{p}^{+}}\perp r^{+}, n_{S_{p}^{-}}\perp r^{-}$. The edge curve of medial surface, also called the fold curve, have the third vectors $\boldsymbol{r}^{\mathbf{0}}$. A $r^{0}$ points from a fold point $m_{0}$ to $S^{0}\in S$. Under these conditions, the boundary surface of the object $S$ is formed from three open manifolds: $S^{+}=m+r^{+}$, $S^{-}=m+r^{-}$, $S^{0}=m+r^{0}$. In medial axis geometry, the vector field $\boldsymbol{r}$ is also a radial map that transport the skeletal geometry to the boundary surface geometry, where Damon derives specific formula in literatures (Damon J et al., 2006). The radial map guarantees 1-1 onto correspondence between the medial surface $m$ and the object boundary surface $S$.

However, important problems exist when digitally modelling the shape of 3D objects extracted from neuroimages by m-reps. To be consistent with the literatures of discrete m-reps, we use the term “spokes” to represent the vectors $\boldsymbol{r}$. The first problem is that since the spokes are required to be perpendicular to the boundary surface, the accuracy of the fold curve is highly required, otherwise the representation of the boundary surface near the fold curves will become sparse. Actually, to describe details, for example a bump, on the boundary surface, the discrete m-rep requires large numbers of spokes on corresponding medial surface. Second, although conditions required to avoid the spokes self-overlap have been proposed in Relaxed Blum representations (Damon J, et al., 2003), the perpendicularity and self-overlap of the spokes are still difficult to satisfy simultaneously on discrete surface. The spokes self-overlap makes it extreme difficult to reconstruct the original shape with high mesh quality.

To improve the descriptions near the fold curves, we define discrete version of $S_{sup}$, $S_{inf}$ and the inscribed medial surface (IMS) as combined sum of discrete surfaces: $S_{sup}=S^{+}+ S^{0}, S_{inf}=S^{-}+ S^{0}, and IMS=m+ S^{0}.$Each point $p$ on IMS has two spokes, the superior spoke $r^{+}$ and the inferior spoke $r^{-}$, that point from $p$ to $S_{p}^{+}$ and $S_{p}^{-}$ respectively. The same to the medial geometry, the superior and inferior spokes satisfy the three properties mentioned above. While, the IMS cancels the third spoke on fold curve, because the fold curve now lives on the boundary surface, making the third spokes become null vectors. By fitting an IMS of the object, the fold curve split the boundary surface of the object into two separate parts, the $S_{sup}$ and $S_{inf}$. Spokes self-overlap remains if we use radial map as the transformation from the IMS to $S_{sup}$ and $S_{inf}$. We notice the method described in (Das SR et al., 2009) that generate surface correspondence by using ﬂow of diﬀeomorphisms can overcome the self-overlap problem, therefore, we refine the original radial map by conformal mapping.

Formally, a diffeomorphism $f:M\to N$ between two surfaces that satisfies $f^{*}ds_{N}^{2}=\lambda ds_{M}^{2}$ is said to be conformal, where $ds_{N}^{2}$ and $ds_{M}^{2}$ are first fundamental forms of surfaces $M$ and $N$. The $f^{*}ds_{N}^{2}$ is the pullback metric induced by $f$. The scaling factor $\lambda$ is called the conformal factor with respect to $f$. The conformal mapping preserves angles and shapes at an infinitesimal scale. This nice property admits to reconstruct the object surface by user-defined parametrization. Also, by overcome the self-overlaps, the conformal mapping improves the mesh quality of the reconstructed boundary surface. The introducing of IMS and conformal mapping refinement avoids strict constraints on fold curves of medial axis while provides adequate descriptions on the boundary surface of the object.

2. **Method and results in Discussion part 1**

The template ARMM of an atlas of hippocampus is used to generate ARMMs of other hippocampi by a deformation-based method, detailed in Method – model evaluation section. To evaluate the method on clinical hippocampus MR images, in this experiment, we implement the method on 7T MRI to test if the deformation field could transform all the elements of ARMM to the target hippocampus.

The point-wise surface correspondence is initially built by SPHARM (Styner M et al., 2006) for better performance of the deformation. We implement the method on five 7T ex-vivo MRI of hippocampi from an open-source study (Adler DH et al., 2018). The demographic information of the five subjects is listed in table S1. The hippocampal surfaces are extracted from the label image by ITK-snap (Yushkevich PA et al., 2006), which are also used as the ground truth to test geometric accuracy of ARMMs by the proposed method. The same hippocampi are also reconstructed by the ds-rep (Liu Z et al., 2021), a medial representation similar to the ARMM while also use a transform-based method to automatically generate population representations for shapes. To fairly compare the performance of the four models, the reconstructed surfaces are setting to contain almost the same number of vertexes (around 1000).

Figure S1 shows an ARMM of a typical 7T hippocampus. The colormap for the left panel of the figure denotes local surface distance between the reshaped surface by ARMM and the ground truth. The most severe surface distance errors are on the medial and lateral of body, with the maximum distance 0.21mm, comparable to the image resolution 0.2mm. The long axis on IMS well captures longitudinal curvature, especially on head, shown in the right panel of figure S1. Also, the transversal axes have been kept nearly orthogonal to the longitudinal axes. Population statistics of the distance errors are listed in table S2. The distance errors are measured by point-wise surface distance Q (detailed in Method – Model evaluation section). The ARMM outperformed the ds-rep on the mean and maximum of the measurement. Also, the reshaped surface by ARMM also has lower standard error of surface distance error than the reconstructed surface by ds-rep. The results indicate that the transformation-based method reaches a resolution-level accuracy (0.2mm) in automatically generation of ARMMs of 7T hippocampi.

Table S1. Demographic information of subjects and MRI resolution.

|  | 7T MRI |
| --- | --- |
| MRI Resolution (mm×mm×mm) | 0.2×0.2×0.2 |
| Group (number of subjects) | CN/AD (3/2) |
| Gender | 1F/4M |
| Age* | 75.8±10.57 |

Abbreviations: F, female; M, male.

*: mean±standard error

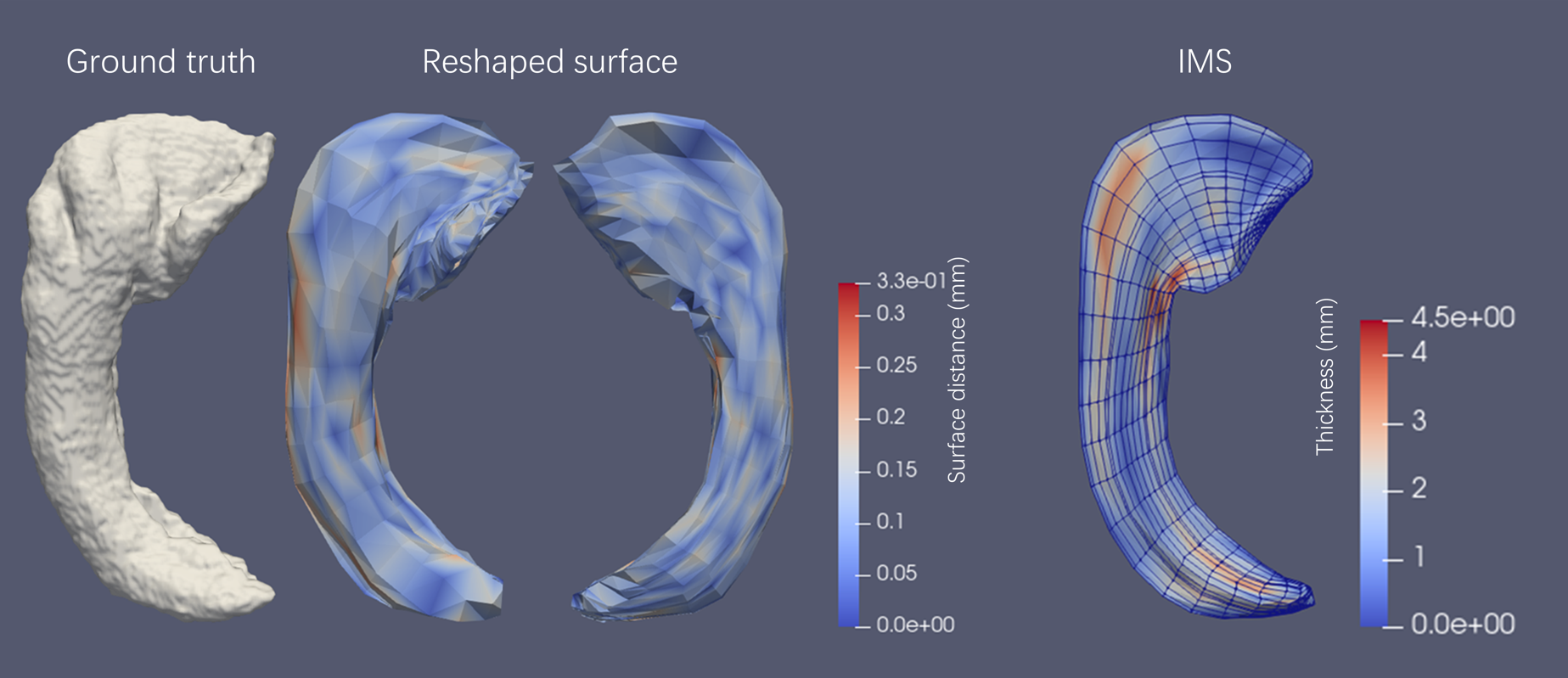

Figure S1. The ARMM of typical 7T hippocampus generated by the deformation-based method. Columns from left to right: the ground truth surface extracted from MRI, the reshaped surface by ARMM, the IMS of hippocampi. A. Surfaces and IMS of a 7T hippocampus.

Abbreviations: IMS, inscribed medial surface.

Table S2. Distance error statistics between the reconstructed surface and the ground truth surface by ARMM and ds-rep respectively.

|  | mean (mm) | standard deviation (mm) | max (mm) |
| --- | --- | --- | --- |
| **7T (ARMM)** | **0.08** | **0.01** | **0.214** |
| 7T (ds-rep) | 0.675 | 0.035 | 0.71 |
